# Utilisation of staphylococcal immune evasion protein Sbi as a novel vaccine adjuvant

**DOI:** 10.1101/413294

**Authors:** Y. Yang, CR. Back, MA. Gräwert, AA. Wahid, H. Denton, R. Kildani, J. Paulin, K. Wörner, W. Kaiser, DI. Svergun, A. Sartbaeva, AG. Watts, KJ. Marchbank, JMH van den Elsen

## Abstract

Co-ligation of the B cell antigen receptor with complement receptor 2 on B-cells via a C3d-opsonised antigen complex significantly lowers the threshold required for B cell activation. Consequently, fusions of antigens with C3d polymers have shown great potential in vaccine design. However, these linear arrays of C3d multimers do not mimic the natural opsonisation of antigens with C3d. Here we investigate the potential of using the unique complement activating characteristics of Staphylococcal immune-evasion protein Sbi to develop a pro-vaccine approach that spontaneously coats antigens with C3 degradation products in a natural way. We show that Sbi rapidly triggers the alternative complement pathway through recruitment of complement regulators, forming a tripartite complex that acts as a competitive antagonist of factor H, resulting in enhanced complement consumption. These functional results are corroborated by the structure of this complement activating Sbi-III-IV:C3d:FHR-1 complex. Finally, we demonstrate that Sbi, fused with *Mycobacterium tuberculosis* antigen Ag85b, causes efficient opsonisation with C3 fragments, thereby enhancing the immune response significantly beyond that of Ag85b alone, providing proof of concept for our pro-vaccine approach.

## Introduction

Opsonisation of an antigen with C3d(g), the final degradation product of complement component C3, results in the co-ligation of the B cell antigen receptor and complement receptor 2 (CR2) on B cells, thereby instigating a profound molecular adjuvant effect, i.e. this co-ligation of receptor complexes lowers the threshold of antigen required for B cell activation by up to 10,000 fold [1-3]. Furthermore, as CR2 is also expressed highly on follicular dendritic cells (FDCs) [4] the presence of C3d(g) on the antigen allows it to be trafficked onto and trapped at the surface of these cells [5]. This provides an essential depot of antigen to support the germinal center reaction and maintain the ongoing immune response including the generation of high affinity antibodies and memory B-cells [3, 4, 6]. B cells can also have an important role as antigen presenting cells (APCs) [7, 8] and have been shown to contribute to T-helper cell priming [9, 10] and therefore, antigen-C3d-CR2 interactions play a key role in humoral immunity [5]. Additionally, C3d activation of T helper cells has also been described in a CR2 independent manner [11], underlining the importance of C3d opsonisation in stimulation of the immune system to respond.

Not surprisingly, this functionality led to the idea that recombinant versions of C3d would make an ideal natural adjuvant and to the subsequent design of linear polymers of human C3d [12]. Indeed, these linear arrays of C3d multimers (3-mer to 20-mer) when fused directly to an antigen can act as potent activators of human B-cells. However, they do not mimic the natural opsonisation of antigens by C3d at a molecular level and do not always enhance immune responses [13]. After activation of C3, C3b attaches directly to the antigen surface via the reactive thioester on the convex face of the protein’s thioester domain (TED). In the presence of complement regulators (factor I (FI) and its co-factors, such as factor H (FH) and CR1) this is rapidly converted to iC3b and then to C3d, exposing the concave CR2 binding site of the TED fragment away from the antigen surface [14]. It is likely that multiple iC3b/C3d molecules attach to complex antigens/pathogen surfaces during the initial activation phases of complement, creating high-avidity binding site for complement fragment receptors.

In the last two decades, structural biology has helped to unveil many of the molecular aspects that are crucial for the activation and regulation of the complement system. Most notable are the crystal structures of the central complement component activation states, native C3 [15], activated C3b and inactive C3c [16]. The structure of C3b in complex with factors B and D [17] subsequently revealed a detailed view of the alternative pathway C3 convertase assembly and its activation, leading to the amplified cleavage of C3 molecules that result in opsonisation and clearance of microbial pathogens and host debris. The covalent attachment of C3b to surfaces does not discriminate between self or non-self surfaces and requires tight regulation to protect host surfaces. Structures of C3b in complex with FH domains 1-4 [18] and domains 19-20 [19, 20] provided insights into protection of host cells [21] and demonstrated how factor H-related proteins (FHRs) function as competitive antagonists of FH, modulating complement activation and providing improved discrimination of self and non-self surfaces [22]. The subsequent structure of the complex of C3b, FH_1-4_ and regulator factor I [23] improved our understanding in the proteolytic cleavage of C3b to the late-stage opsonins iC3b or C3dg and provided the basis for the regulator-dependent differences in processing and immune recognition of opsonized material.

Here we investigate the potential to harness the unique complement-stimulatory characteristics of *Staphylococcus aureus* immunomodulator Sbi to develop ‘pro-vaccines’. Sbi components would trigger natural complement activation in the host and coat antigen surfaces with complement component C3 degradation products, thereby enhancing the degree of immunogenicity of target antigens. Research from our lab previously revealed that Sbi contains two domains (III and IV), which bind to the central complement component C3 and cause futile fluid phase consumption of this component [24]. Therefore, these two domains of Sbi offer the potential to not only coat an antigen with the natural adjuvant C3d, but also to generate anaphylatoxins and the full range of C3 opsonins. Such an approach has the clear potential to activate many immune cells unlike recombinant C3d fragment-based adjuvants of the past, that, due to the restricted expression pattern of CR2, were largely focused to B cells. Furthermore, the direct activation of complement close to the target antigen (with the associated anaphylatoxin generation) may be critical for generating appropriate inflammatory immune responses, both humoral and cellular; needed to immunise against complex pathogenic targets.

In this study we first investigate the molecular mode of action of Sbi-III-IV and evaluate the importance of the tripartite complex formation between Sbi, C3d and complement regulators factor H (FH) or factor H-related proteins (FHRs) for complement activation. Based on these findings, we then tested whether our pro-vaccine strategy would be successful by using *Mycobacterium tuberculosis* antigen 85b (Ag85b) as a model antigen in a fusion construct containing Sbi domains III and IV. We show that this Sbi-Ag85b conjugate is opsonized by C3 degradation products in serum, and when administered to mice, leads to an enhanced immune response *in vivo*, but only in mice that possess C3 and complement receptor 1 and 2, demonstrating proof of concept for this adjuvant compound.

## Results

### Sbi-III-IV triggers C3 consumption via activation of the alternative complement pathway, forming a covalent adduct with C3b

To investigate the molecular details of the C3 futile consumption caused by Sbi, a protein construct consisting of domains III and IV (Sbi-III-IV) was incubated with normal human serum (NHS) and analyzed using western blotting. As seen previously [24], we found that Sbi-III-IV-induced C3 consumption results in the deposition of metastable C3b molecules onto serum proteins, causing the formation of high molecular weight C3b covalent adduct species with serum proteins (Figure 1a). Immuno-blotting analyses using a polyclonal anti-Sbi antibody (Figure 1b) reveals that a small fraction of Sbi-III-IV molecule also forms a covalent adduct with a nascent C3b molecule that is subsequently converted into a smaller Sbi-iC3b adduct as a result of proteolytic processing by serum proteases. In addition, we show that Sbi-III-IV-induced C3 consumption coincides with the release of the C3a anaphylatoxin fragment (Figure 1c), and the proteolytic activation of factor B (FB) (Figure 1d), confirming the alternative complement pathway as the driving force behind this process. Pre-incubation of serum with Sbi-III-IV results in the loss of serum hemolytic ability caused by the futile consumption of fluid C3 (Figure 1e). Without pre-incubation, the Sbi-III-IV construct does not protect rabbit erythrocytes from lysis in NHS under AP conditions.

**Figure 1.**
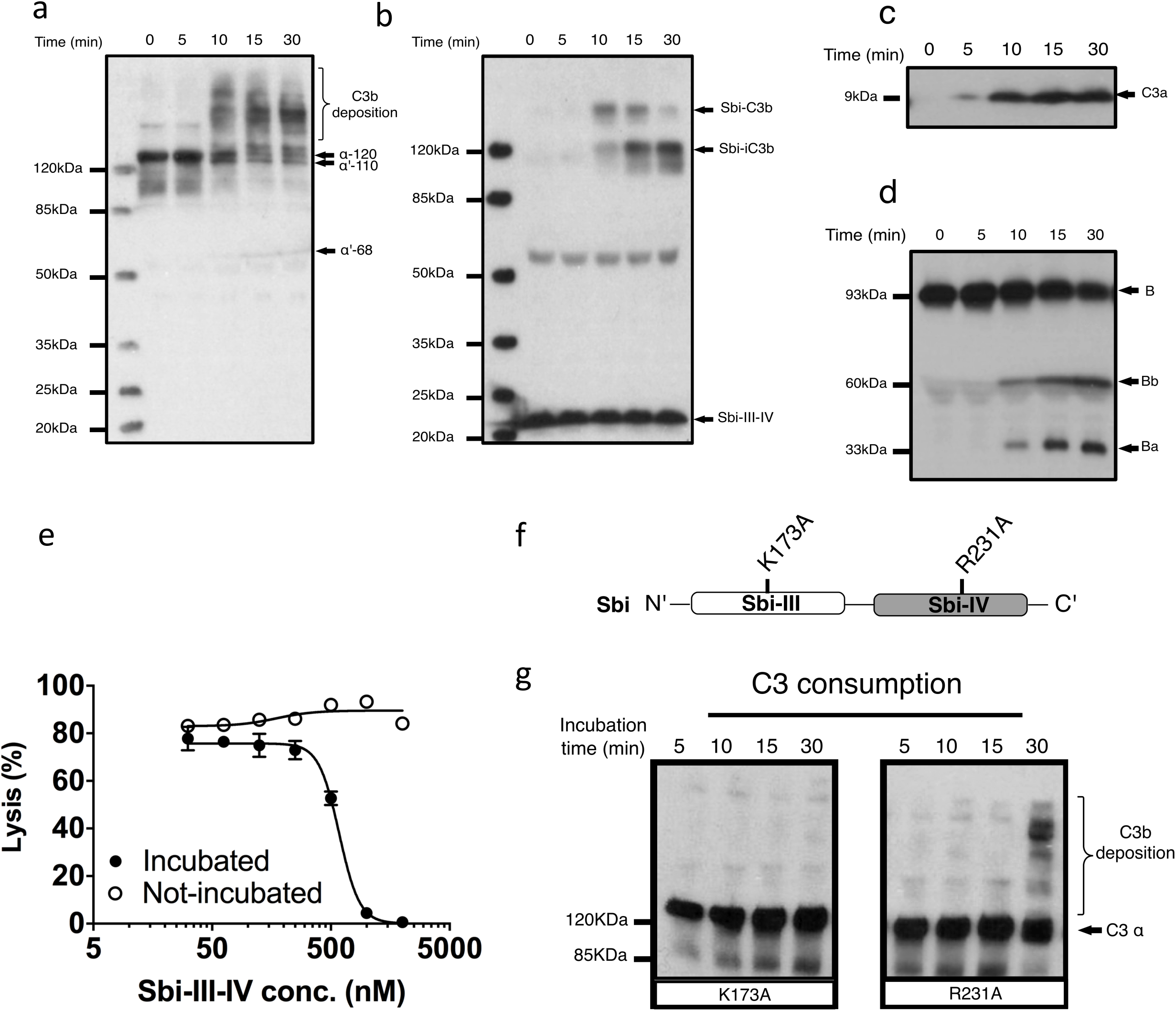
Sbi-III-IV induces C3 futile consumption via the alternative complement pathway and thereby causes C3b adduct formation and C3a anaphylatoxin production. (a) C3 activation and C3b deposition in NHS after incubation with 10 μM Sbi-III-IV, visualized using anti-C3d western blot analysis. C3b adducts formed with serum proteins are indicated. Positions of α-120 (C3) α’110 (C3b) and α’-68 (iC3b) are indicated. The 10 minute lag-time in C3 activation we observe in the presence of excess Sbi-III-IV (10 μM) correlates with the delay reported in the natural C3 “tick-over” process, required for supplying the critical enzymatic component for the initial fluid phase Alternative Pathway (AP) C3 convertase. (b) Sbi-C3b adduct formation, visualized with anti-Sbi western blot. These adducts migrate at higher than expected molecular weights (Sbi-α’110: ∼160 kDa and Sbi-α’68: ∼120 kDa, with expected molecular weights of 125 and 83 kDa, respectively) which is caused by the high pI of the Sbi-III-IV construct (pI = 9.3). Sbi-III-IV has a molecular weight of 14.8 kDa, but migrates to ∼22 kDa in SDS-polyacrylamide gel due to the positively charged electrophoresis buffer. (c) C3a anaphylatoxin production, followed using anti-C3a western blot analysis (showing only the low molecular weight region). (d) FB cleavage, monitored by anti-FB western blot analysis. (e) Concentration dependent Sbi-III-IV induced C3 consumption, studied by a rabbit erythrocytes haemolytic assay. Rabbit erythrocytes were exposed to normal human serum pre-incubated with Sbi-III-IV (incubated, closed circles) and normal human serum with Sbi-III-IV added at the start of the experiment (not incubated, open circles). (f) Schematic representation of the relative positions of point mutations that display the most profound functional defects, K173A and R231A. (g) C3 consumption profiles of Sbi-III-IV mutants K173A and R231A. For b-e and h, one representative blot of three independent experiments was shown. For (e), four independent measurements of two experiments were shown. The mean and SD for each measurement were calculated for all datasets. Curves were fitted using non-linear variable slope (four parameters) function in GraphPad Prism.

### Sbi domain III residue K173 is essential for complement consumption

In order to gain understanding of the individual roles of Sbi domains III and IV in AP activation, a systematic site-directed mutagenesis approach was used, mainly focusing on charged and polar amino acids (for details see Supplementary Table S1 and Figure S1). Functional screening of these mutants identified K173, located within Sbi domain III (Figure 1f), as an essential contributor to triggering C3 consumption. Sbi mutant K173A shows no complement activation after 30 minutes incubation with human serum, demonstrating a comparable complement activation defect to the previously identified C3d binding mutant R231A [24, 25] (Figure 1f and g), located in Sbi domain IV. Assessment of the C3d binding affinity, using *switch* SENSE (Table 1 and Supplementary Figure S2A), shows that contrary to R231A the C3d binding capacity of K173A is unaffected, indicating it is essential for the role for domain III in the futile consumption of C3. Interestingly, our *switch* SENSE analyses of the C3d binding characteristics also shows a reduced hydrodynamic diameter for K173A compared to WT and the C3d impaired binding mutant R231A, indication that this mutation in domain III results in a more compact Sbi:C3d complex (Table 2 and Supplementary Figure S2B). A more detailed structural analysis of these conformational changes follows below.

**Table 1.** Sbi-III-IV:C3d interacKon affinity determined by *Switch* SENSE.

| | $K_d$ (nM) | $k_{ON}$ ($M^{-1}S^{-1}$) | $k_{OFF}$ ($S^{-1}$) |
| --- | --- | --- | --- |
| WT:C3d | 5.0±0.8 | 5.9±1.0×10<sup>5</sup> | 3.0±0.1×10<sup>-3</sup> |
| K173A:C3d | 5.8±1.2 | 5.3±0.9×10<sup>5</sup> | 3.0±0.4×10<sup>-3</sup> |
| R231:C3d | No binding |  |  |

**Table 2.** Sbi-III-IV:C3d complex hydrodynamic diameter determined by *Switch* SENSE.

| | $D_H$ (nm) of Sbi-III-IV | $D_H$ (nm) of Sbi:C3d complex |
| --- | --- | --- |
| WT | 4.3±0.1 | 6.6±0.2 |
| K173A | 3.8±0.3 | 5.0±0.3 |
| R231A | 4.7±0.3 | No binding |

### Sbi-III-IV enhances binding of FH or FHRs to C3 breakdown products

In a previous study, we reported that Sbi-III-IV binds C3 isoforms in combination with the C-terminal part of FH (FH_19-20_), forming tripartite complexes [26]. Many FHRs share SCR modules with high FH_19-20_ sequence identity [22] particularly FHR1 which has been demonstrated to have significant complement dysregulation potential [21]. Thus, we investigated the potential role for Sbi-III-IV in mediating the formation of tripartite complexes with C3 fragments and FHR-1, FHR-2 or FHR-5.

On a C3b opsonised surface plasmon resonance (SPR) sensor chip, the presence of wild-type Sbi-III-IV clearly enhanced the binding of FH, FHR-1, FHR-2, FHR-5 as well as FH_19-20_ (at fixed concentrations of 100, 12.5, 20, 25 and 20nM, respectively) to the surface in a concentration dependent manner (Figure 2a). However, in the case of the K173A mutant, tripartite complex formation with FH or FHR-1, 2, 5 or FH_19-20_ is significantly impaired, showing decreased binding and more rapid dissociation compared to WT Sbi-III-IV (Figure 2b and 2c). We also co-injected Sbi-III-IV with FH or FHR-1, flowing opsonized iC3b or amine-coupled C3d(g) across the surface. On these surfaces, the fold-changes in FH (or FHR-1) binding levels were also enhanced even at reduced Sbi-III-IV concentration (Supplementary Figures S3A-D).

**Figure 2.**
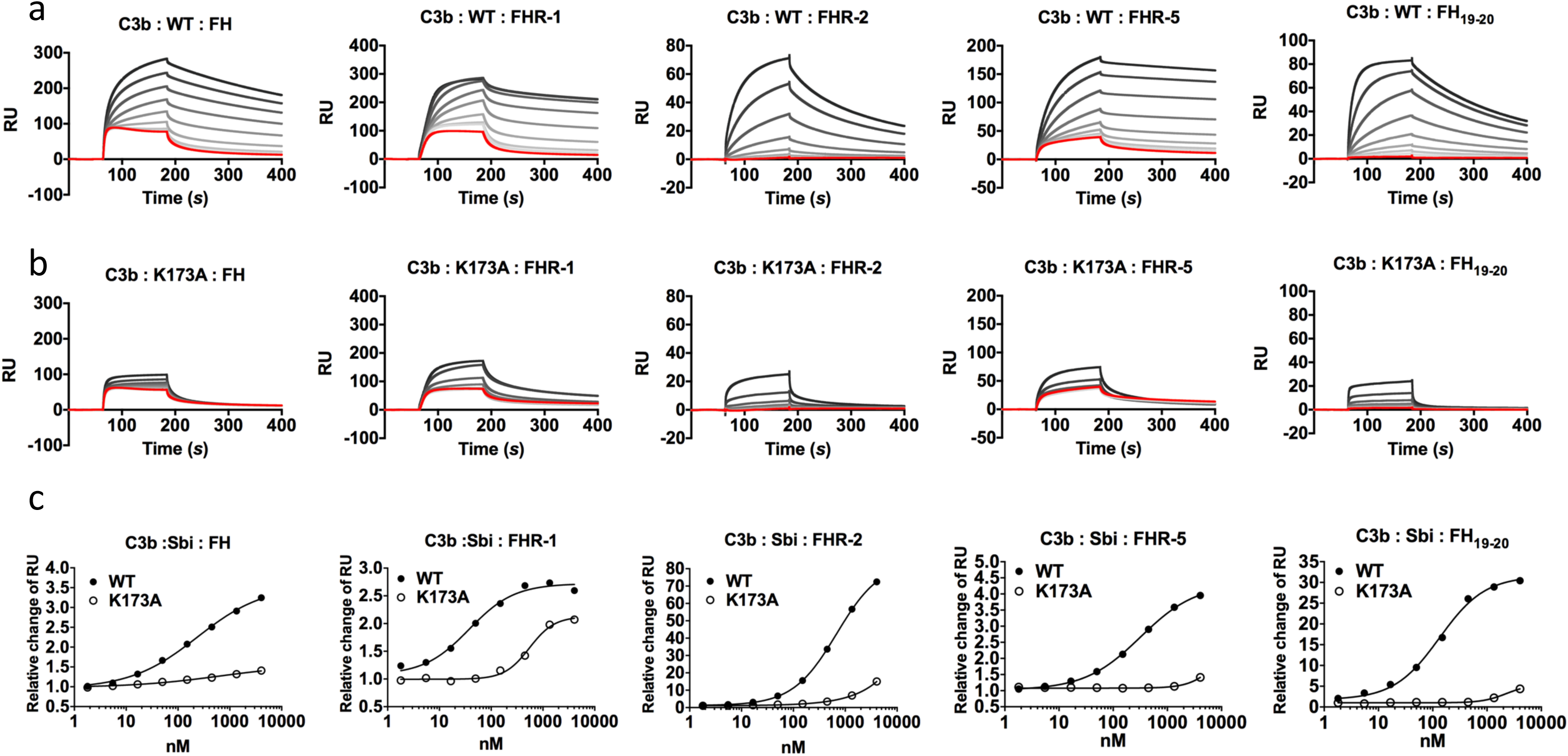
Surface plasmon resonance analyses of tripartite complexes. A triply diluted concentration series (4050 to 1.8 nM) of (a) WT or (b) K173A Sbi-III-IV were co-injected with plasma purified FH, recombinant FHR-1, FHR-2, FHR-5 or FH_19-20_. The red response curves were indicative of binding experiment in the absence of Sbi. The co-injection experiments of a fixed analyte concentration in combination with increasing Sbi concentration were depicted by increasingly dark lines. (c) Relative changes of Sbi-III-IV mediated FH (or FHR) binding to C3b. By subtracting the co-injection sensorgram (i.e. Sbi+FH) with the corresponding Sbi binding dataset (Supplementary Figure S3), the changes in FH (or FHRs) binding was deduced (Supplementary Figure S3). Changes in FH (or FHRs) binding were expressed as the relative change, derived from dividing the Sbi mediated binding by the FH (or FHR) only control, using the response-difference values at the equilibrated binding point (173.5 *s*). Each sensorgram is representative of two experiments. Relative change curves were fitted using non-linear variable slope (four parameters) function in GraphPad Prism.

### Sbi-III-IV acts as a competitive antagonist of FH via the recruitment of FHRs

Our SPR data, described above, show that Sbi-III-IV enables FH or FHR-1, 2 and 5 binding to the C3 activation fragment C3b and late-stage proteolytic fragments iC3b and C3d(g). To further our understanding of the mechanism of FH or FHR recruitment and the contribution of these tripartite complexes to AP complement activation, we used a rabbit erythrocyte haemolytic assay. In the presence of Sbi-III-IV and endogenous FH (and FHRs), in NHS, addition of recombinant FHR-1 or FHR-2 resulted in significantly enhanced C3 consumption (Figure 3a), as evidenced by the reduction in erythrocyte lysis in a concentration dependent manner. In the absence of Sbi-III-IV only baseline C3 consumption was observed. Although FHR-5 alone can reduce erythrocyte lysis in a concentration dependent manner, as described previously [27], in the presence of Sbi-III-IV C3 consumption by FHR-5 is clearly enhanced (Figure 3b). As predicted, the results in Figure 3c indicate that the observed reduction in erythrocyte lysis caused by C3 fluid phase consumption, in the case of FHR1 and likely the remaining FHRs, is mediated by the C-terminal SCR domains of the protein rather than the N-terminal domains.

**Figure 3.**
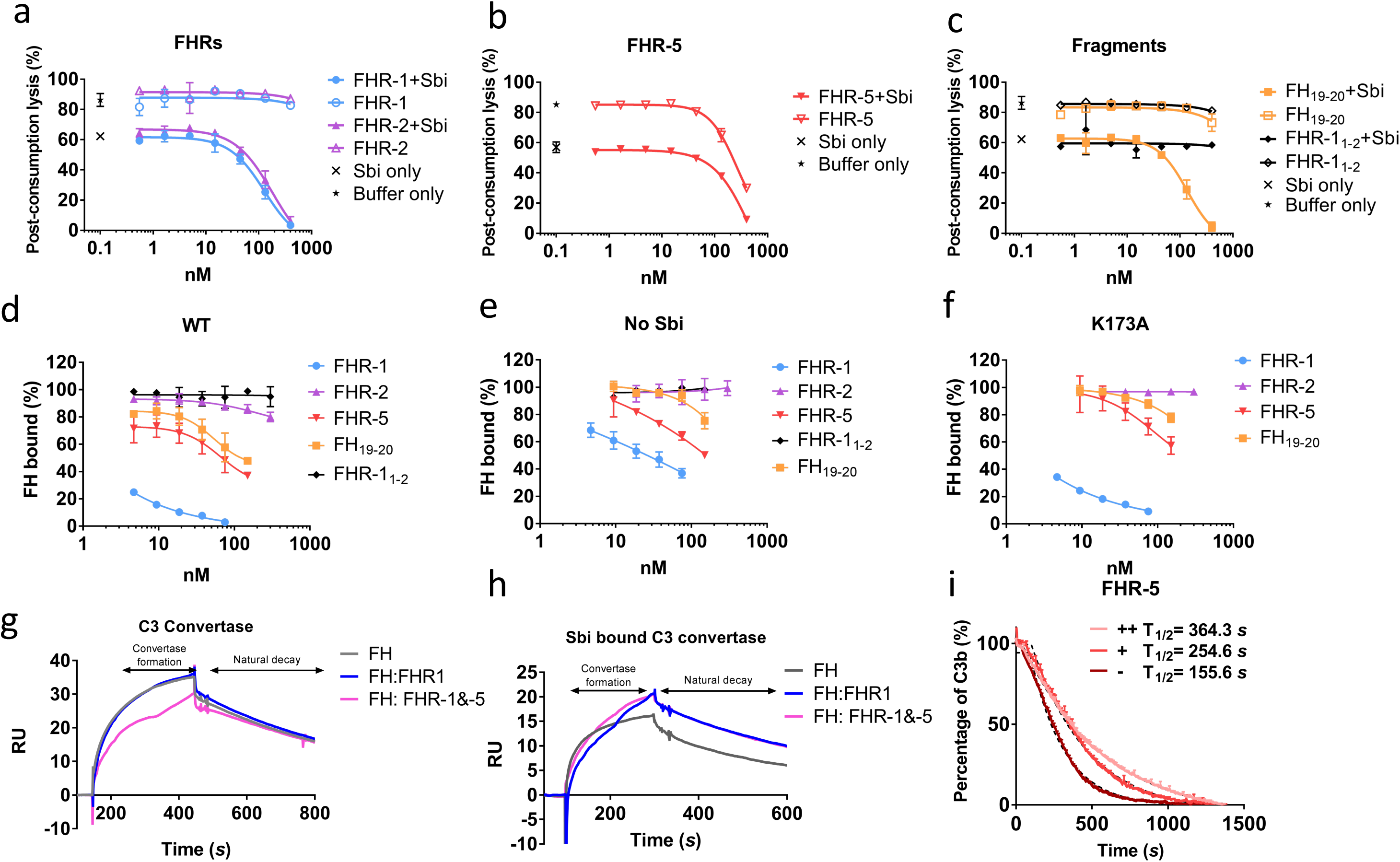
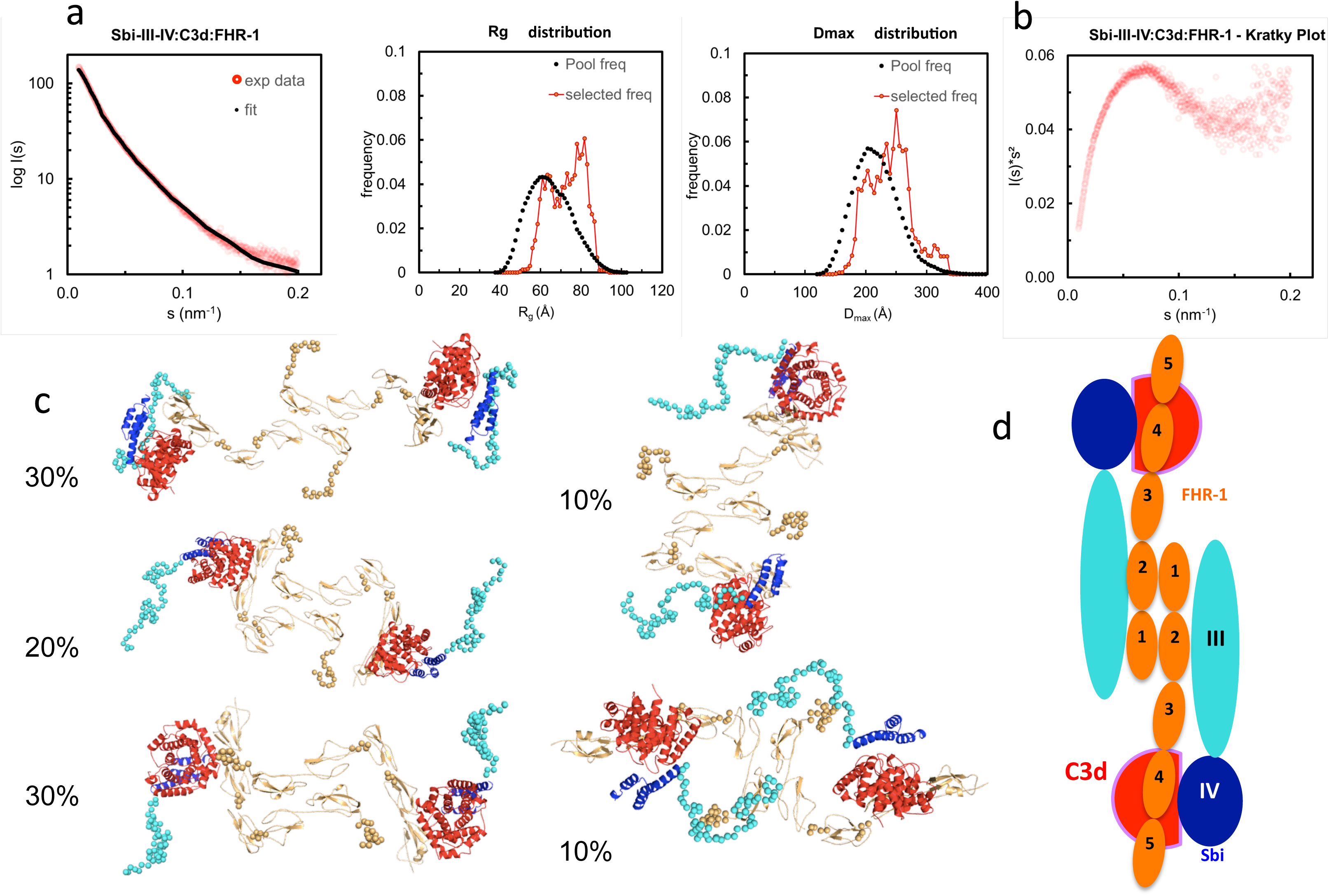
Functional characterization of tripartite complexes in complement AP regulation. NHS was incubated with Sbi-III-IV, in combination with specified reagents or just buffer, the consumption of AP activity was indicated by the protection of rabbit red blood cell from lysis. (a) Pre-incubation of recombinant FHR-1 or −2 with the presence or absence of Sbi-III-IV. (b) Pre-incubation of recombinant FHR-5 with the presence or absence of Sbi-III-IV. (c) Pre-incubation of recombinant FH_19-20_ or FHR-1_1-2_ with the presence or absence of Sbi-III-IV. Using an ELISA assay, the ability of FHR-1, −2, −5, FH_19-20_ or FHR-1_1-2_ to modulate FH binding to a C3b coated surface was studied in the presence of WT (d), no (e) or K173A Sbi-III-IV (f). C3 convertase formation in the absence (g) or presence of Sbi-III-IV (h) was assessed by flowing factor B (500 nM) and factor D (100 nM) in the presence of FH (2000 nM) or FH +FHR-1 (2000 and 200 nM) or FH+FHR-1&-5 (2000, 200 and 20 nM) across a surface amine coupled with 500 RU C3b. To form Sbi bound C3 convertase, experiments were conducted in addition of 2000 nM of Sbi-III-IV. Detailed experimental and data processing procedures are provided in *material and methods* and Supplementary Figure S3E. (i) Percentage of intact C3b derived from continuous recording of ANS fluorescence changes between 465-475 nm spectrum. Baseline C3b breakdown curve (-) was recorded in the presence of FH and FI, interference caused by the addition of FHR-5 (+) or FHR-5 in combination of Sbi (++) was also examined. The data for FHR-1 and FHR-2 are presented in Supplementary Figure 3F. Normalized data was depicted in solid lines, simulated breakdown curves were shown as dotted-lines. Each curve represents the mean value of three independent experiments. For (a-f), the mean and standard deviation for each measurement was calculated; For (g-h), each sensorgram is representative of two experiments. For (i), simulated breakdown curves were fitted using one phase exponential decay function in GraphPad Prism.

Whilst our SPR and rabbit erythrocyte assay clearly indicate that *in vitro* Sbi-III-IV can recruit FHRs in tripartite complexes with C3b and thereby enhance fluid phase complement consumption, it has to be taken into account that the physiological molar concentrations of FHR-1, 2 and 5 are 13-164 fold less than that of FH [21, 28]. To further investigate the potential competitive binding between FH and the FHRs in Sbi-III-IV mediated tripartite complexes, we used an ELISA-based assay where we applied FH (25 nM) and Sbi-III-IV (1 μM, in the presence of a concentration range of FHRs (9.3 – 150nM)) onto a C3b coated plate. Subsequently, we assessed the percentage of FH bound using monoclonal antibody OX-24. Figure 3d shows that FHR-1 can compete with FH to bind C3b, decreasing the percentage of residual FH bound to C3b from ∼70% at the lowest FHR-1 concentration to ∼30% at the highest concentration. In the presence of Sbi-III-IV WT this effect is dramatically increased with only ∼25% residual FH bound at the lowest FHR-1 concentration, reducing to ∼0% at the highest FHR-1 concentration (Figure 3e). These results clearly indicate that Sbi-III-IV can preferentially recruit FHR-1 to form a tripartite complex with C3b. Similarly, enhancement of recruitment was observed with FHR-5 and fragment FH_19-20_ but only weakly with FHR-2. Although unable to activate complement, Sbi-III-IV mutant K173A is still able to compete for the binding of FHR-1 in the presence of FH, but its ability to enhance binding of FHR-2 and FHR-5 to C3b is clearly affected (Figure 3f).

To assess the potential AP de-regulatory roles of the Sbi-III-IV mediated tripartite complexes. We subjected them to a novel C3 convertase decay acceleration activity (DAA) assay and a fluid phase C3b co-factor activity (CFA) assay [29, 30]. We demonstrated that in absence of Sbi, FHR-1 failed to antagonise FH efficiently and show a difference in the level of C3 convertase formation (Figure 3g). Co-injection of FHR-1 and −5 shows reduced C3 convertase formation, which is in accordance with the results from a previous study [31]. However, the presence of Sbi (2μM) potentiates the FH antagonising effect of FHR-1, and to a lesser extent that of FHR-5, at a physiologically relevant concentration ratio (FH 2000 nM: FHR-1 200 nM: FHR-5 20 nM), resulting in increased C3 convertase formation on a C3b surface (Figure 3h). The baseline C3b breakdown rate was acquired in the presence of FH (0.160μM) and FI (0.017μM), and subsequent measurements were performed in the presence of FHR alone (0.32 μM) and in combination with Sbi-III-IV (1 μM). As shown in Figure 3i and Supplementary Figure S3F, the presence of FHR-1, 2 or 5 increases the C3b fluid phase half-life to different degrees, with FHR-5 showing the largest increase in half-life. Most interestingly, the C3b half-life could be further extended by the addition of Sbi-III-IV.

### The Sbi-III-IV:C3d:FHR-1 tripartite complex forms a dimer in solution

To investigate the structural characteristics of the Sbi-III-IV:C3d:FHR1 tripartite complex, we used small angle X-ray scattering (SAXS). The scattering profile collected at an equimolar mixing ratio is shown in Figure 4 in log plot (a) as well as Kratky plot (b). The featureless descend in the log plot and the plateau in the latter is characteristic for scattering of particles that are, at least partially, disordered.

**Figure 4.**
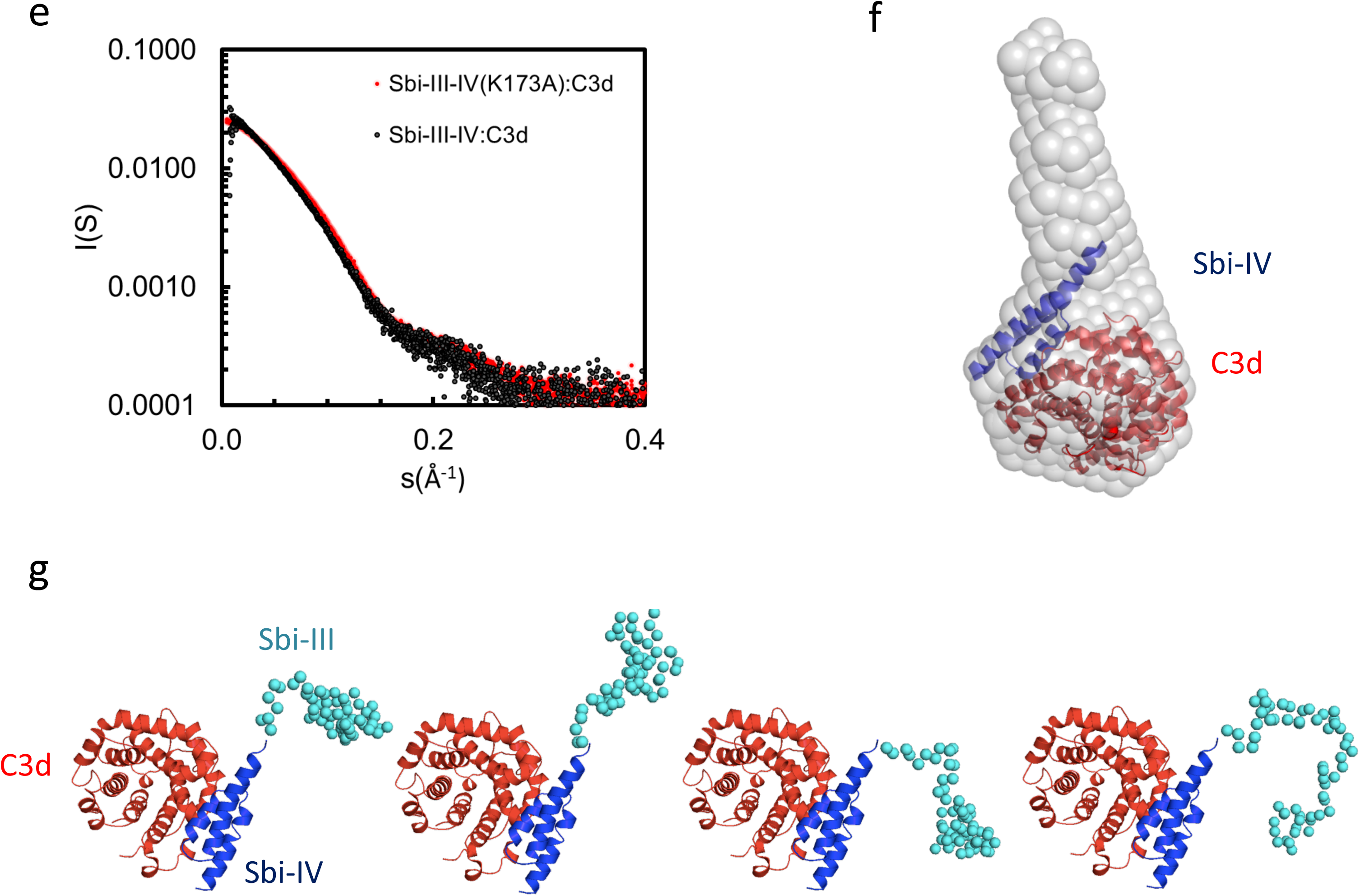
Structural analysis of the Sbi-III-IV:C3d:FHR-1 tripartite complex. SAXS solution structure analysis and EOM modeling of the Sbi-III-IV:C3d:FHR-1 tripartite complex: (a) Left panel, fit of the selected ensemble of conformers to the experimental scattering. Radius of gyration (*Rg*, middle panel), particle maximum dimension (*Dmax*, right panel), and distribution histograms of the selected conformers versus the pool. (b) Kratky plot of the tripartite complex. c) Examples of rigid body models of the selected conformers corresponding to the histogram peaks. The volume fraction of each species is indicated. The relative positions of C3d, Sbi-III-IV and FHR-1 in the dimeric tripartite complex are indicated, with C3d in red, Sbi-IV in dark blue, Sbi-III in turquoise and FHR-1 in orange. (d) Schematic representation of the dimeric Sbi-III-IV:C3d:FHR-1 tripartite complex. (e) Comparison of the solutions structure of wild-type Sbi-III-IV:C3d and mutated version Sbi-III-IV(K173A):C3d of the dual complex. Radius of gyration (*Rg*), particle maximum dimension (*Dmax*), and distribution histograms of the selected conformers versus the pool are shown in Supplementary Figure S4A. (f) *Ab initio* shape reconstruction shown as gray spheres in comparison to the partial crystal structure Sbi-IV:C3d (2wy8). (g) Examples of rigid body models. Complete set of models as well as flexibility assessment is presented in supplement Figures S5. C3d in shown red, Sbi-IV in dark blue, and Sbi-III in turquoise.

The SAXS data and the overall parameters obtained (Supplementary Table S2) suggest that the complex is largely dimeric but rather flexible in solution. Quantitative flexibility analysis was performed using the ensemble optimisation method EOM [32], which fits the experimental data using scattering computed from conformational ensembles. Models with randomized linkers were generated based on the known structures of FHR-1_1-2_ (3zd2, [21]); FH_18-20_ (3sw0, [33]), containing the equivalent of FHR-1_3_; FH_19-20_:C3d complex (2xqw, [20]), corresponding with FHR-1_4-5_; and the Sbi-IV:C3d complex (2wy8, [34]). To account for the dimerisation, P2 symmetry was applied, using the FHR-1_1-2_ dimer interface as seen in the crystal structure (3zd2). The distributions of the overall parameters in the selected structures compared with those of the original pool (Figure 4c) suggests that the complex is rather flexible with a slight preference for extended structures in solution. The subset of most typical models (and the volume percentage of their contribution) shown in Figure 4c indicate that in addition to the expected contact sites with C3d, Sbi-III domain appears to also interact with FHR-1, corroborating the functional results described above. Figure 4d shows a schematic representation of the dimeric Sbi-III-IV:C3d:FHR-1 complex observed in solution.

### K173A restricts the conformational freedom of Sbi domain III

To examine the possible structural effects of the K173A substitution in Sbi domain III, SAXS data was collected on the Sbi-III-IV(K173A):C3d complex and compared to the wild-type Sbi-III-IV:C3d complex published previously [34]. The experimental scattering pattern collected at 240 µM (∼12 mg/ml) is presented in Figure 4e and the structural parameters derived from this data are given in Supplementary Table S2. The estimated molecular mass (MM) of the solute agrees within the errors with the values predicted for a 1:1 complex (∼15 kDa + 35 kDa). At lower concentration a decrease in the MM estimates is observed which suggests that the complex slowly begins to dissociate. The previously described wild-type Sbi-III-IV:C3d data on the other hand, suggests that at higher concentrations, higher oligomeric species are present, thus, for the comparison here, data collected at 0.6 mg/ml is shown. The faster descend of the wild-type data, which translates to a larger Rg, suggests that rearrangements of the flexible N-terminus lead to a more elongated particle (Rg wild-type= 32.8 Å) as compared to K173A mutant (Rg K173A = 30.6 Å). This is in strong agreement with the *switch* SENSE analysis of the C3d binding characteristics, which show a reduced hydrodynamic diameter for K173A compared to WT (Table 2).

The *ab initio* low resolution models of the complex reconstructed from the highest concentration data using DAMMIF [35] showed a large cone shaped molecule with a volume of 124 nm^3^ (Figure 4f). The resolution of the reconstruction is estimated to be 29 +/- 2 Å [36]. The large base of the cone can accommodate the crystal structure of Sbi-IV:C3d complex (2wy8) [34]. The extra space at the tip of the cone would be sufficient to harbor the 60 N-terminal residues comprising the Sbi-III domain. A more detailed modeling was conducted with the program Coral [37], utilizing the available high-resolution model of Sbi-IV:C3d and allowing for 60 additional beads to be added that mimic the missing Sbi-III domain. Twenty independent Coral runs were performed which all yielded models with a more or less structured N-terminal region, suggesting that Sbi-III-IV(K173A) in complex with C3d is conformationally restricted compared to wild-type Sbi-III-IV. This is further supported by the narrow distributions obtained with EOM (Supplementary Figure S4B). Surprisingly, whilst repeating these analyses using a proposed alternative binding mode of the Sbi-IV:C3d complex (represented by 2wy7), where Sbi-IV is seen bound at the convex face of C3d, the χ^2^ value is greatly improved (Supplementary Figure S4C). With this modeling approach a similar restricted conformation is observed for the N-terminus of Sbi-III-IV(K173A). Further studies are currently being conducted to further investigate the potential physiological relevance of this alternative Sbi-IV:C3d binding mode.

### A fusion construct of Sbi-III-IV with M. tuberculosis Ag85b activates the AP

To test the potential of Sbi-III-IV to induce C3d opsonisation in a vaccine setting, a recombinant construct was designed whereby Sbi-III-IV is fused to *Mycobacterium* protein Ag85b (Figure 5a, and detailed in Supplementary Figure S5A). Based on the SAXS structure of the Sbi-III-IV:C3d:FHR-1 tripartite complex, revealing the importance of a flexible and extended conformation of Sbi domain III, we decided to attach Ag85b at the C-terminus of Sbi domain IV and included a long flexible linker between Sbi-IV and Ag85b to ensure accessibility and flexibility of the functional domains. Expressed and purified fusion protein was subsequently structurally and functionally characterized.

**Figure 5.**
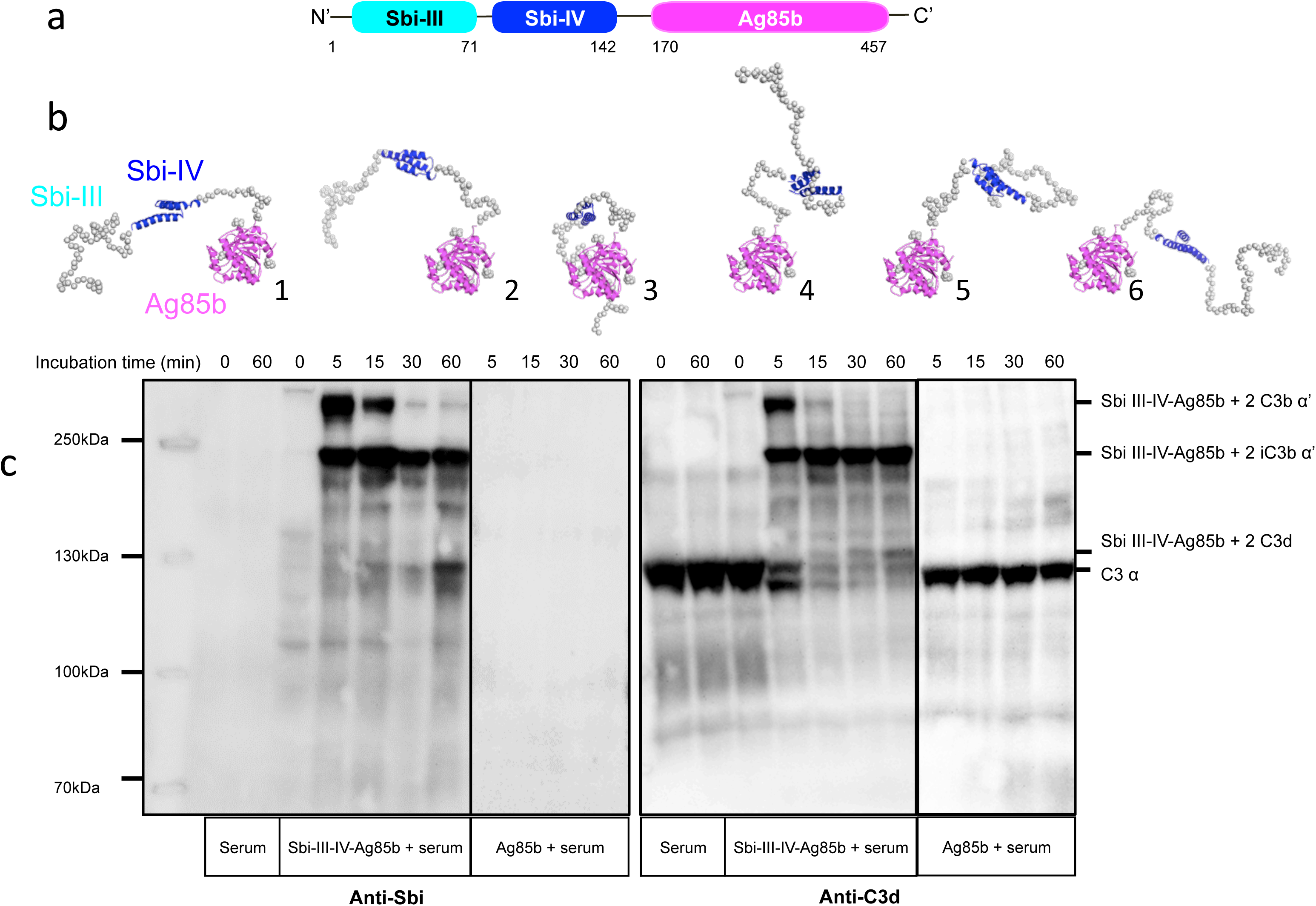
Structural and functional analysis of the Sbi-III-IV-Ag85b fusion protein. (a) Schematic structure of the Sbi-III-IV-Ag85b fusion protein (see fo details Supplementary Figure S9). (b) SAXS analysis of the fusion protein indicates a monomeric molecule with a radius of gyration of R_g_=3.7 nm and maximum particle size of D_max_ =15 nm. The various molecular mass estimation range from 44 −51 kDa and are comparable with a predicted monomer mass of 50 kDa. The 10 independent *ab initio* models obtained with DAMMIF are similar to each other, and according to the χ^2^ values that estimate the goodness of the fit, the final structures fit well with the experiment. More detailed modelling with Coral and EOM show that the flexibility of the missing structural information is restricted. (c) C3 activation and C3-fragment deposition in NHS after incubation with Sbi-III-IV-Ag85b (100 μM) or Ag85b (100 μM), visualized using anti-Sbi and anti-C3d western blot analysis. Resultant higher molecular weight bands with Sbi III-IV-Ag85b were identified as Sbi-III-IV-Ag85b with two covalently attached C3b α’ chains; Sbi-III-IV-Ag85b with two iC3b α’-68 chains and Sbi-III-IV-Ag85b with two C3d molecules. Ag85b alone is unable to activate C3 as indicated by the presence of an intact C3 α-chain.

Circular dichroism analysis of the Sbi-III-IV-Ag85b fusion indicates that the protein construct is folded and that the secondary structural elements of both parent proteins have been preserved (Supplementary Figure S5B). SAXS data obtained for the fusion protein demonstrate that both the Ag85b domain as well as Sbi-IV domain are accessible (Figure 5b and Supplementary Figure S5C).

Functional activity of the Sbi-III-IV-Ag85b fusion construct was assessed using an AP complement activity assays (WIESLAB, Euro Diagnostica), showing strong C3 depletion activity (Supplementary Figure S5D), whilst Ag85b on its own showed no complement activating properties. These results confirm that the complement activating properties of Sbi III-IV are not impaired as part of the fusion construct. The western blot analyses presented in Figure 5c and Supplementary Figure S5E confirm these results, showing both C3 activation and opsonisation by the Sbi-III-IV-Ag85b fusion construct when incubated with NHS. Interestingly, C3 activation and consumption occur more rapidly with the fusion construct when compared to Sbi-III-IV (Figure 1a and 1b) under the same conditions. Whilst Sbi-III-IV shows opsonisation with a single molecule of C3b (Figure 1b), the Sbi-III-IV-Ag85b fusion is opsonized by 2 molecules of C3b that over time degrade to iC3b and C3d (Figure 5c and Supplementary Figure S5E). Interestingly, opsonisation of Ag85b with C3 fragments also occurs when co-incubated with Sbi-III-IV in NHS.

### Sbi-III-IV acts as an adjuvant in mice when immunized with Ag85b

Based on the ability of Sbi-III-IV to activate complement (Figures 1b, 1c (human serum) and 6a (mouse serum)) and opsonise Ag85b with complement C3 break down fragments (Figure 5c), we expected that this new fusion protein when injected into mice would elicit a greater immune response to the Sbi-III-IV-Ag85b fusion protein than Ag85b administered alone (in PBS). Indeed, wild-type C57bl/6 mice immunized I.P. (or I.V., data not shown) with Sbi-III-IV-Ag85b generated a greater than 4 fold increase in immune response initially and following the boost when compared to Ag85b alone (Figure 6b). Furthermore, when mice were immunized with a mixture of Sbi-III-IV and Ag85b (not fused together), this also resulted in a significantly improved immune response, corroborating the role of C3 fragment opsonisation of the antigen in this process (see Supplementary Figure S5E). Subsequent, studies using C3^-/-^ and *Cr2*^-/-^ mice clearly demonstrated that C3 and C3 breakdown fragment receptors (CR1 and CR2) were essential for this “adjuvant” function, respectively (Figure 6c). Overall, these data clearly suggest that complement AP dysregulation function of the Sbi-III-IV domain can be harnessed to improve immune responses through the coating of antigens with C3 breakdown fragments.

**Figure 6.**
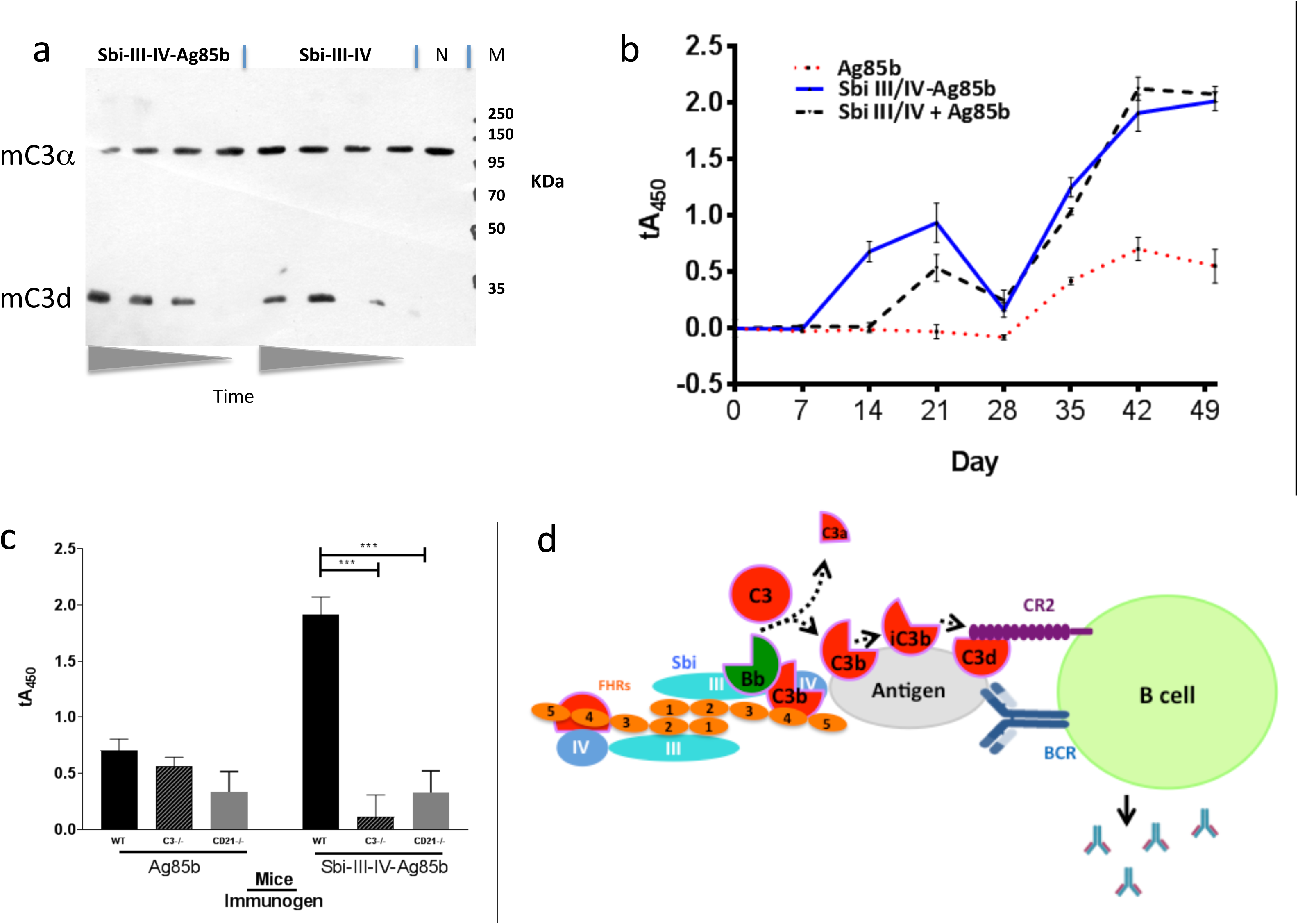
Sbi-III-IV is an effective adjuvant in mice. (a) Freshly prepared CD21^-/-^ mouse serum was mixed with Sbi-III-IV-Ag85b or just Sbi-III-IV. The reaction was stopped at various time points (0, 30, 60, 120 minutes). Western blot was developed with rabbit anti-C3 at 1/1000 and goat anti-rabbit at 1/2000. C3d is shown as confirmation that C3 has been activated and broken down. (N) is *Cr2*^-/-^ serum incubated for 120 minutes with saline. (b) C57Bl/6 mice (groups of 6) where immunised intraperitoneally with either 2.7µg Sbi-III-IV-Ag85b protein, 2µg Ag85b, or 0.7µg Sbi-III-IV plus 2µg Ag85b in 150mM NaCl solution, followed by weekly bleed and boosted (day 28) before terminal bleed at day 49. Serum IgG reactivity to Ag85b was measured over time by ELISA. Sera was diluted 1/50 and the mean absorbance ± SEM of each mouse group is shown. All data has been normalised to the day 0 average of all WT mice. (c) The previous experiment was repeated in C57Bl/6 mice deficient of C3 (C3^-/-^)and complement receptor type I and 2 (*Cr2*^-/-^). Data is representative of at least 2 repeats. (d) Schematic representation of the dimeric Sbi-III-IV:C3d:FHR-1 solution structure providing a nidus for AP C3 convertase generation that overwhelms local complement regulators, leading to the opsonisation of the nearby antigen surface by C3 break-down products that help facilitate the co-ligation of the B cell antigen receptor (BCR) with complement receptor 2 (CR2) thereby lowering the threshold for B cell activation.

## Discussion

Previous work from our group [24] revealed that *Staphylococcus aureus* immunomodulator Sbi binds complement component C3 within the thioester domain of C3 or the C3dg portion of the molecule and resulted in futile consumption of C3 via uncontrolled activation of the AP. In this study, we endeavored to both understand the mechanism of action of Sbi-III-IV and harness it; in order to develop pro-vaccines which would trigger natural complement activation and thereby coat antigen surfaces with complement component C3 degradation products, generate anaphylatoxins at the site of immunisation and strongly enhance the immunogenicity of antigens (Figure 7d).

The seminal studies by Pepys *et al* [38], using C3 activating/depleting agents including cobra venom factor and Zymosan, clearly demonstrated intact C3 function was important for the T-dependent response [38]. A molecular mechanism explaining this effect was established by Fearon and colleagues [39] supported by studies in both C3 [40] and *Cr2* (complement receptor type I and II) knock-out mice [41]. Fearon *et al* exploited these findings and established that multiple copies of C3d, in a linear trimer, could enhance antigen-specific responses up to 10,000 fold [3]. However, the initial potential of trimeric C3d, as a highly potent molecular adjuvant, has not been realized and the reason(s) for this remain(s) unclear. One possible explanation is that the artificial linear trimer structure fails to represent naturally opsonised antigen, and consequently does not provide sufficient CR cross-linking or additional inflammatory signals for the B cell (or APC) activation threshold to be reached. One possible approach to overcome this is attaching more C3d to test antigens, but that approach is also limited [42]. In the light of these and other findings [11, 13, 43], we considered that with understanding of the mode of action of Sbi-III-IV we might be able to develop a new complement activation based immune adjuvant.

The first clue to a mechanism for Sbi’s ability to rapidly activate the AP came from monitoring Sbi-III-IV treated NHS in a time course using anti-C3 and anti-Sbi immuno-blotting. Here, we demonstrated that metastable C3b not only attaches covalently to serum proteins but also to Sbi-III-IV itself; as a transacylation target (Figure 1). This makes sense in the respect that Sbi’s affinity to C3 obviously places it in close proximity to the site of complement turnover and we speculate that C3b deposited on Sbi-III-IV could help extend the fluid phase half-life of C3b, preventing FH and FI from binding and inactivating as normal, perhaps similar to covalent adducts of C3b with IgG [44, 45].

However, as we have shown previously, Sbi-III-IV also interacts with complement regulators FH and FHR-1, in addition to binding C3b and its degradation products, thereby forming tripartite complexes [26]. We next investigated whether Sbi-III-IV acts as a competitive antagonist of FH via the recruitment of FHR-1 and 5 into tripartite complexes and that FHR-1 can effectively displace FH from the tripartite complex. To this end, data from our systematic site-directed mutagenesis screen brought to light several Sbi mutants with complement activation defects (Figure 1, Supplementary Figure S1). For instance, we demonstrated that an alanine substitution in Sbi domain III at position 173 resulted in a dramatic reduction in C3 consumption activity (Figure 1g). Notably, although a similar effect was observed with a previously identified mutation in domain IV with impaired C3d binding (R231A), K173A showed only slightly impaired C3d binding capacity (Table 1) suggesting a different mechanism. We therefore postulated that the K173A mutant would be ideal to elucidate the structural and functional role of Sbi domain III in the activation of complement and found that K173 in Sbi domain III was crucial for the recruitment of FHR-5 and that the K173A mutation only slightly affects FHR-1 binding (Figures 2 and 3). These findings implicate a direct role of Sbi domain III in the tripartite complex formation with these FHRs and that this likely occurs via interactions with the C-terminal SCR domains that share sequence identity with FH_19-20_. We confirm this by showing that increasing concentrations of recombinant FHR-1, FHR-2, FHR-5 and FH_19-20_ in serum indeed potentiate Sbi-III-IV mediated C3 consumption, whilst the N-terminal SCRs (FHR-1_1-2_) fails to do this (Figure 3).

We also observed that Sbi greatly enhances the binding of FHRs to C3b, thereby antagonizing FH activity, as shown by the C3 convertase decay accelerating activity (DAA) assay (Figure 3 g and h). These results imply that the FHR-1 or FHR-5 containing tripartite complexes can protect the AP C3 convertase, aiding the consumption of C3. These findings further enhance the notion that the FHR family has diversified AP de-regulatory functions, where FHR-1 seems more efficient in counteracting the DAA of FH, whilst in contrast FHR-5 potently antagonizes the cofactor activity (CA) of FH. The observed Sbi-III-IV mediated shift in the complement regulatory balance towards C3 activation could potentially be further enhanced by the formation of homo/heterodimeric forms of FHR-1 with itself and with other FHRs (FHR-2 and FHR-5) [12]. These data link to an ongoing evolutionary ‘arms race’ where FH was initially hijacked by *S. aureus* to protect it from complement [46] and then FHRs (devoid of intrinsic complement regulatory activity) were evolved/deployed by the host to compete with FH on that surface and restore complement opsonisation of the pathogen [22]. Perhaps the release/secretion of Sbi from *S. aureus* is a more recent event in this arms race with the host, which takes the C3b/C3 convertase binding potential away from the bacterial surface and leads to local rapid fluid phase consumption of complement, i.e. local decomplementation and bacterial survival/propagation. Our understanding of the role and complexity of FHRs in immune evasion strategies is still in its infancy [46], but this study underlines the potency of another strategy in this process.

Using FHR-1 as a ‘model’ dimerization domain containing FHR, structural analysis of the Sbi-III-IV:C3d:FHR-1 tripartite complex, using SAXS, indeed suggests the formation of a dimer mediated by FHR-1 domain 1 and 2 and provides details of the role of the extended unfolded nature of domain III in the binding of FHR-1 (Figure 4). The molecular basis of the preferential binding of FHR-1 over FH cannot easily be explained on the basis of differences in amino acid sequence between the two complement regulators, since their C3d binding regions (SCR 4-5 of FHR-1 and SCR 19-20 of FH) share 99% sequence identity. However, our SAXS analyses, and binding studies using C3d(g) or iC3b as ligands (Figures 3 and 4), indicate that the C-terminal regions of FHR proteins are readily exposed, unlike those of FH that exist in a “latent” conformation with the C-terminal part of the protein folded back and partially blocked [47-50]. The dimeric physiological state of FHR-1 and the other FHRs tested in this study is also likely to enhance their ability, due to increased avidity, to assemble a tripartite complex.

Analysis of the hydrodynamic volume of the Sbi-III-IV:C3d complex using *switch* SENSE highlighted a significant contraction of the normally extended conformation Sbi-III-IV structure [34] caused by the K173A substitution in domain III (Table 2). SAXS analysis confirms these findings, showing a partially kinked N-terminal structure of domain III in K173A with reduced conformational freedom (Figure 4e-g). The contraction of the Sbi-III-IV structure caused by the K173A substitution suggests that the normally flexible and extended conformation of domain III plays an important role in the recruitment of FHRs, especially FHR-5 into the tripartite complex after the initial interaction between Sbi-IV and C3b. Previous structural analyses of the Sbi’s domain III, using NMR, revealed that this domain is indeed natively unfolded [51].

Based on the structural and functional information described here we decided to construct a Sbi-III-IV-Ag85b fusion construct that could be used to test its effect on the immune response against this model antigen *in vivo*. We chose *Mycobacterium tuberculosis* Ag85b, a fibronectin-binding protein with mycolyltransferase activity [52], because it is known to be immunogenic and previously suggested as a vaccine candidate [53]. Indeed, there is evidence that Ag85b can elicit both humoral and cellular immune reactions in patients with TB, but there is conflicting evidence of its efficacy as a vaccine [54, 55], suggesting adjuvants may improve its overall immunogenicity. This target also gives scope to allow further testing in animal models of disease [56]. Structural analysis, using Circular Dichroism and SAXS confirmed that secondary structural elements of both parent proteins have been preserved in the fusion protein construct and that the crucial functional Sbi domains are accessible for interactions with complement (Figure 5 and Supporting Figure S5). We also show that the Sbi-III-IV-Ag85b fusion construct can induce AP activation and is opsonized with C3 breakdown products (Figure 5c).

With AP activation in human and mouse serum confirmed (Figures 5 and 6), we opted to use straightforward immune response, IgG titre, analysis to demonstrate the potential of Sbi-III-IV to trigger complement *in vivo* and act as a vaccine adjuvant in a mouse model, in a similar manner to many previous studies [57]. Our data herein firstly indicates that Sbi-III-IV can activate mouse complement in an analogous manner to that of the human complement system. This obviously allows direct analysis of these pro-vaccine compounds in both mouse and human model systems (a huge advantage to previous C3d based adjuvants) [13], indeed Sbi-III-IV has acted as a C3 activator in all species tested thus far (data not shown). As predicted from the *in vitro* work, the opsonisation of fusion proteins or co-immunised antigen by mouse complement breakdown fragments results in a significant increase in the immunogenicity of Ag85b, with increased IgG titres noted in the presence of fused or co-immunised Ag85b (Figure 6). The adjuvant function both increased the intensity of the response and the rate of the response when compared to Ag85b immunized alone. We will need to further explore the potency of this response to that of common adjuvants and with a mix of target antigens to fully assess the utility of Sbi-III-IV as a universal vaccine adjuvant. For instance, comparison of the action of Sbi-III-IV to the Glaxo-Smith-Kline’s adjuvant systems, particularly AS01 [58], or to MF59 [59] may be of key interest and recent approaches may provide ideal pre-clinical model systems to facilitate this [60, 61] before progression to clinical studies. The work is ongoing but the data herein demonstrate the initial proof of concept.

In summary, we have demonstrated that Sbi-III-IV triggers consumption of complement component C3 via activation of the alternative complement pathway, by acting as a competitive antagonist of FH via the recruitment of FHRs into dimeric tripartite complexes that can protect C3b bound to Sbi (Figure 6d). It is likely this provides a stable nidus for alternative pathway mediated C3 convertase generation, i.e. local fluid phase C3bBb generation that overwhelms any local complement regulators, providing the potential for bystander lysis or opsonisation of surfaces. Our ability to harness this potential, targeting complement opsonisation to the surface of an antigen (in this case from *Mycobacterium)* and therefore use Sbi-III-IV as a vaccine adjuvant clearly demonstrates Sbi-III-IV has great potential for use with a range of antigens across multiple species, including humans, although more work remains to make that a reality.

## Materials and Methods

### Proteins, antibodies and sera

Factor H (FH), C3b, factor B (FB), factor D (FD), factor I (FI), properdin (FP), FI-depleted serum, goat anti-human C3 polyserum and goat anti-human FB polyserum were purchased from Complement Technologies (Tyler, TX). FHR-1_1-2_, FHR-1, −2 and −5 used in the tripartite complex reconstruction and binding competition assay were produced using Chinese Hamster ovary cell culture (as previously described [62]). Horse radish peroxidase (HRP)-conjugated rabbit anti-goat immunoglobulin polyserum and HRP-conjugated Streptavidin were acquired from Sigma Aldrich. HRP-conjugated goat anti-rabbit immunoglobulin G (Thermo Fisher, catalog no. 815-968-0747), HRP-conjugated rabbit anti-mouse immunoglobulin G (Thermo Fisher, catalog no. 31452) and biotin-conjugated FH monoclonal antibody OX24 (catalog no. MA5-17735) were purchased from Thermo Fisher Scientific. The goat anti-human FH polyclonal serum (catalog no. 341276-1ml) that was previously used to detect human FH and FHR-1 was purchased from Merck Millipore. Human C3 was purified from mixed pool citrated human plasma (TCS Bioscience, PR100) using polyethylene glycol 4000 precipitation, anion and cation exchange chromatography as previously described [63]. A pET15b-C3d construct was acquired from Prof. David E. Isenman and transformed into *Escherichia coli* (*E. coli*) stain BL21 (DE3), recombinant C3d was then expressed and purified using a previously described protocol [64]. Lyophilized normal human serum (NHS) was purchased from Euro Diagnostica (catalog no. PC300). Additional proteins and antibodies are described in the specific experimental sections.

### Sbi-III-IV constructs

The expression and purification of the N-terminally 6×His tagged recombinant Sbi-III-IV from a pQE30:*sbi-III-IV* construct were described previously [24].

### Sbi-III-IV mutagenesis

Mutations in the Sbi-III-IV sequence were introduced using the QuikChange II XL site-directed mutagenesis kit (Agilent Technologies), the primers used are listed in Supplementary Table S1. The mutated pQE30:*sbi-III-IV* plasmids were sequenced to confirm the success of the mutagenesis. SDS-PAGE profiles of all the Sbi-III-IV mutant proteins used in this study are shown in Supplementary Figure S1.

### Sbi-III-IV induced C3 consumption assay

Lyophilized NHS was re-suspended in chilled dH_2_O to a 2× concentration. Equal volumes of 2×NHS and Sbi (10 μM) were combined. Sbi treated sera were then incubated in a thermocycler at 37°C for 30 min. Treated serum samples were collected at time intervals, 0.5 μl of serum was loaded on an SDS-PAGE gel analyzed under reducing condition. The proteins were Western blotted, and the blots were probed with anti-C3d, anti-Sbi, anti-C3a or anti-factor B antibodies.

A hemolytic assay was modified from a previously published procedure [65] to measure Sbi induced consumption of C3. Briefly, rabbit erythrocytes (TCS Bioscience) were resuspended in GVB buffer (5 mM veronal, 145 mM NaCl, 10 μM EDTA, 0.1 % (w/v) gelatin) by washing three times via centrifugation at 600 *g* for 6 mins. The concentration of rabbit red cells to be used in each experiment was determined by adding a stock of 5 μl of erythrocytes to 245 μl of water to give complete lysis and then re-adjusting cell concentration until an optical density reading of 0.7 (A_405_) was reached. Lysis experiments were conducted in two steps, first, 15 μl of NHS, 5 μl of Mg+-EGTA (70 mM MgCl_2_ and 100 mM EGTA), 20 μl of protein in E2 buffer was mixed and pre-incubated at 37 °C for 30 mins. Subsequently, 5 μl of rabbit erythrocyte was added and incubated for an additional 30 mins at 37 °C. At the end of the incubation, 150 μl of quenching buffer (GVB supplemented with 10 mM EDTA) was added. The cells were pelleted by centrifugation at 1500*g* for 10 min, and absorbance (A_405_) of 100 μl of supernatant measured. Post-consumption lysis percentage was calculated as 100×((A_405_ test sample-A_405_ 0% control)/(A_405_ 100%-A_405_ 0% control)).

### In vitro complement activation using Sbi III/IV-Ag85b in mouse serum

Mouse serum was collected from male *Cr2*^-/-^ mice by cardiac puncture and allowed to clot fully on ice for 4 hours followed by separation of serum by centrifugation at 2000g in a refrigerated centrifuge. Serum was then mixed with Sbi III/IV or Sbi III/IV-Ag85b, ensuring that the amount of Sbi III/IV in each preparation was equivalent. The reaction was stopped at 0, 30, 60 and 120 minutes, by the addition of reducing sample buffer, boiled for 5 min and analysed on a 10% SDS-PAGE gel. After transfer to nitrocellulose the blots were probed with Rabbit anti-C3d (1/1000, DAKO, A0063) and Goat anti-Rabbit-HRPO (1/2000, 111-035-046-JIR, Stratech), developed with ECL substrate (Pierce) and exposed to X-Ray film for 2 min.

### switchSENSE kinetic analysis

A switchSENSE DRX 2400 instrument (Dynamic Biosensors) was used to characterize the binding kinetics and protein size changes based on *switch* SENSE technology [66, 67]. Purified Sbi-III-IV-cys, K173A, R231A and their ligand C3d were sent to Dynamic Biosensor’s protein analyzing facility for binding kinetic and hydrodynamic diameter analysis. In the case of a protein binding event, based on the real-time measurements of the switching dynamics in a range of ligand concentrations, binding rate constants (*k*_ON_ and *k*_OFF_) and dissociation constants (*K*_D_) can be analysed [67]. Alternatively, under saturated binding conditions, the switching dynamic of the protein (or protein complex) can be compared with the switching dynamics of bare DNA and with a biophysical model with which the size of the immobilized protein (or protein complex) can be determined. For determination of Sbi-III-IV:C3d binding kinetic parameters, 130 nM, 100 nM, 70 nM and 40 nM of C3d were applied sequentially onto the Sbi-III-IV immobilized microchip. All Sbi:C3d complexes’ hydrodynamic diameters were estimated at a C3d concentration of 130 nM.

### Fluorometric assay

Fluorometric C3b breakdown assay was performed using a black 96 well microplate (Thermofisher, M33089) in a TECAN Spark 20M temperature-controlled fluorescence plate reader. Excitation was at 386 nm and emission was recorded at 475 nm with a 20-nm bandwidth. The control C3b breakdown rate, performed in PBS, contained 100 µl of 1 µM C3b, 160 nM FH, 17 nM of FI and 10 µM ANS, and was scanned every 5 *s* for 15 min. To study the interruption of C3b breakdown, 32 nM of FHR was either added alone or in combination with 1 µM of Sbi-III-IV. Data were collected at 25°C, normalized by Excel using the equation “Percentage of C3b=((*F*_X_-(*F*_15min_))/(*F*_15min_-*F*_0min_))*100” and plotted by Graphpad Prism.

### Small angle X-ray scattering

Synchrotron radiation X-ray scattering from solutions of the Sbi-III-IV:C3d:FHR-1 tripartite complex, the Sbi-III-IV(K173A):C3d complex, and the Sbi-III-IV-Ag85b fusion protein were collected at the EMBL P12 beamline of the storage ring PETRA III (DESY, Hamburg, Germany). Images were collected using a photon counting Pilatus-2M detector and a sample to detector distance of 3.1 m and a wavelength (λ) of 0.12 nm covering the range of momentum transfer (s) 0.1 < s< 4.5 nm^-1^; with s=4πsinθ/λ. Different solute concentrations were measured using a continuous flow cell capillary. To monitor radiation damage, 20 successive 50 ms exposures were compared and frames displaying significant alterations were discarded. The data were normalized to intensity of the transmitted beam and radially averaged; the scattering of the buffer was subtracted, and the different curves were scaled for solute concentration. The forward scattering I(0), the radius of gyration (Rg) along with the probability distribution of the particle (p(r)) and the maximal dimension (D_max_) were computed using the automated SAXS data analysis pipeline SASFLOW [68].

For the Sbi-III-IV-Ag85b fusion protein data quality was improved with SEC-SAXS mode and the parallel analysis of light scattering data in a similar manner as described in Gräwert *et al*. [69]. Frames comprising solely the monomeric version of the fusion protein were averaged and used for further processing after background subtraction.

The molecular masses (MM) were evaluated by comparison of the forward scattering with that from a reference solution of BSA and based on the Porod volumes of the constructs. With SAXS, the former estimation of MM is within an error of 10%, provided the sample and standard concentrate are determined accurately. DAMMIF was used to compute the *ab initio* shape models. For this, 10 independent models fitting the experimental scattering curves were generated and compared to each other. More detailed modelling was obtained with Coral. Here, existing partial crystal structure of the Sbi-IV:C3d complex was extended with 60 additional beads placed at the N-terminus of Sbi-IV to mimic the missing Sbi-III domain. Here too, 10 independent runs were performed, and the degree of variation addressed. Further analysis of the flexibility of the samples was addressed with Ensemble Optimization Method (EOM). For this, ensembles of models with variable conformations are selected from a pool of randomly generated models such that the scattering from the ensemble fits the experimental data, and the distributions of the overall parameters (e.g. D_max_) in the selected pool are compared to the original pool.

The proteins in the Sbi-III-IV:C3d:FHR-1 tripartite complex were combined 1:1:1 at a concentration of 45 µM. The Sbi-III-IV(K173A):C3d complex were formed at a 1:1 ratio at 240 µM (12 mg/ml). PDB structure 2wy8 (Sbi-IV:C3d complex) was used to model the complex using and compared with SAXS data previously recorded [34]. The Sbi-III-IV-Ag85b fusion protein was provided at 29, 72 and 145 µM concentrations (1.45, 3.6 and 7.2 mg/ml, respectively). The samples were dialysed against PBS, which was also used for background subtraction. From all samples concentration series were measured to exclude any concentration dependent alterations.

### Surface Plasmon Resonance

Tripartite complexes were analyzed by surface plasmon resonance (SPR) technology using a Biacore S200 (GE Healthcare). All experiments were conducted at 25°C on CM5 chips, using HBST (10 mM HEPES, 150 mM NaCl, and 0.005% Tween 20, pH 7.4) as running buffer, which was optionally supplemented with 1mM of MgCl_2_ (HBST+) to be compatible with AP amplification condition. On the chip surface 800 RU of C3b was opsonized via AP C3 convertase through a method described before [70, 71]. The iC3b surface was produced by injecting of repetitive cycles of FH and FI across a C3b opsonized surface, the completeness of the conversion was confirmed by the inability of FB binding. A separate chip surface was made by amine coupling 600 RU of recombinant C3d (CompTech, USA). In all SPR experiments, response differences were derived using the signal from a flow cell to subtract the parallel reading from a reference flow cell that blocked with carbodiimide, *N*-hydroxysuccinimide and ethanolamine. Analytes were injected in duplicate (at 30 μl/min for 200 *s*) followed by running buffer for 300 *s* and a regeneration phase involving injection of regeneration buffer (10 mM sodium acetate, 1 M NaCl pH 4.0) for 60 *s*. To analyze Sbi-III-IV binding and the assembly of tripartite complex, concentration series of Sbi-III-IV WT or K173A were flowed cross separately or co-injected with FH, FHR-1, FHR-2, FHR-5 or FH_19-20_ at a fixed concentration (100, 12.5, 20, 25 or 20 nM respectively).

The C3 convertase DAA assay was performed on a CM5 chip amine coupled with 500 RU C3b, using HBST^+^ as running buffer throughout. A mixture of analytes for building C3 convertase were flowed across, including FB and FD in addition to various FH reagent combinations (FH or FH and FHR-1 or FH, FHR-1 and −5). The various FH reagents combinations were also flow across separately in order to derive the sensorgram for C3 convertase. To examining Sbi bound C3 convertase, 2 µM of wild-type Sbi-III-IV was added to the mixture of analytes for building C3 convertase. The various FH reagent combinations spiked with Sbi were flowed across separately in order to derive the sensorgram for Sbi bound C3 convertase. Each injection cycle includes Injection of the C3 convertase mixture for 200-300 *s*, followed by running buffer for 300-400 *s* and two consecutive 60 *s* regeneration phases.

### FH/FHR-1 Competition assay

C3b was diluted in carbonate buffer (pH 9.5) and coated on to wells of a Nunc MaxiSorp plate (0.25 µg/well) for 16 h at 4°C. The wells were blocked with PBST (PBS with 0.1% Tween 20) supplemented with and 2% BSA for 1 h at 37 °C, and then washed with PBST buffer. Doubly diluted concentration series (9-600 nM) of FHRs, FH_19-20_, FHR-1_1-2_ in PBST-2% BSA were then added to the wells, together with a constant concentration of FH (25 nM) and Sbi-III-IV (1000 nM). The plate was incubated for 1 h at 37 °C, then washed with PBST. 50 µl of monoclonal anti-FH antibody OX-24 (specific to the FH SCR domain 5) diluted with PBS-2% BSA (0.6 µg/ml) was added to the wells and the plate incubated for a further 1 h. The wells were washed with PBST, and 50 µl sheep anti-mouse IgG (1:5000 dilution in PBST-2% BSA) was added to the wells for 1 h at 37 °C. The wells were washed again and the conjugate was detected using TMB ELISA substrate solution, which was added to the wells for 5 min. The colour reaction was stopped by 10% H_2_SO_4_ and the plate was read at *A*_450_ using a plate reader.

### Design and purification of the Sbi-III-IV-Ag85b fusion construct

The DNA sequence coding for Sbi-III-IV (*sbi*_448-798_) was fused to the 5’ end of the DNA sequence for Ag85b (*ag85b*_121-975_) via a linker region of 84 bp (Supplementary Figure S6). The fusion gene was commercially synthesized and ligated into the pET15b vector, containing an ampicillin resistance cassette and a T7 promoter. The pET15b:*sbi-III-IV-ag85b* plasmid was verified using sequencing, and the resulting construct encoded an N-terminally his-tagged Sbi-III-IV-Ag85b protein. *E. coli* BL21 (DE3) cells harbouring the pET15b:*sbi-III-IV-ag85b* plasmid were grown in LB broth supplemented with 100 µg/ml ampicillin to an *A*_600_ = 0.4-0.6. Protein expression was induced with 0.5 mM IPTG and by incubating the cells at 17°C for 16 h. Bacteria were harvested, lysed using sonication (80% amplitude, for six 10 s bursts) in the presence of protease inhibitor cocktail (set VII-Calbiochem, Merck), and the protein initially purified using nickel-affinity chromatography (His-Trap column, GE Healthcare) with a gradient of 0-0.5 M imidazole in 50 mM Tris, 150 mM NaCl, pH 7.4. It was further purified using size-exclusion chromatography (Hi-Load 16/60 Superdex S200 column, GE Healthcare) equilibrated in 20 mM Tris, 150 mM NaCl, pH 7.4. Fractions containing protein were pooled and concentrated. Protein concentration was measured at *A*_280_.

### Analysis of Sbi-III-IV-Ag85b fusion protein AP complement activity

Alternative pathway (AP) activity of Sbi III-IV-Ag85b-treated NHS samples was analysed using the ELISA-based WIESLAB^®^ (Euro Diagnostica) complement system AP assay. Sbi-III-IV-Ag85b was mixed with normal human serum (NHS) at a 1:1 volume ratio and incubated for 30 min at 37 °C in a thermal cycler. Treated serum was then diluted with AP diluent (blocking the activation of the other two complement pathways) by 1 in 20. From this point the manufacturer’s instructions were followed. A blank (AP diluent), positive control (NHS) and negative control (heat-inactivated NHS) were also recorded. Complement activation was converted to residual AP activity (%) using the equation: (sample - negative control)/(positive control - negative control) x 100.

### C3 fragment deposition on Sbi-III-IV-Ag85b fusion protein

The method used is similar to that described for WT Sbi-III-IV. Lyophilised NHS (Euro Diagnostica) was re-suspended in chilled dH_2_O. Sbi-III-IV-Ag85b (100 μM) was mixed with NHS in a 1:1 ratio, and incubated for 1 h at 37 °C in a thermocycler. Samples were taken at regular intervals (0, 5, 15, 30, 60 min), and separated by SDS-PAGE followed by Western blot analysis using either rabbit anti-Sbi (1.5: 5000 dilution), rabbit anti-C3d (1.5:5000 dilution) or mouse anti-Ag85b (1:1000 dilution) polyclonal antibodies and detected using HRP-conjugated secondary antibodies (1:2500 goat anti rabbit or 1:1000 goat anti mouse). NHS-only was used as a negative control.

### Measurement of immune response to Sbi-III-IV-Ag85b fusion protein

Eight week old male mice (wild-type C57bl/6, C3^-/-^ and *Cr2*^-/-^) were bled by tail vein venesection at day −2. Mice were then immunized at day 0 with molar equivalent doses of Ag85b alone (2μg), Sbi-III-IV-Ag85b (fusion protein), Sbi-III-IV alone or a mixture of Sbi-III-IV and Ag85b, as appropriate. Mice were then bled weekly thereafter and plasma stored at −80°C until required for batch analysis. Mice were boosted at day 28 and sacrificed at day 42.

For analysis of IgG response to Ag85b by ELISA, 96 well plates (NUNC maxisorb) were coated with 1 µg/ml Ag85b (Abcam, UK) or 1.35 µg/ml Sbi III/IV-Ag85b in carbonate buffer at 50 µl per well and incubated at 4°C for 16 h. Plates were washed with 0.01% PBS-Tween and a 1% BSA blocking solution was applied for 1 h at 20°C. Serum samples were diluted to 1/50 or 1/100 in 0.01% PBS-Tween, added at 50 µl per well and incubated for 1 h at 20°C. Plates were washed and secondary antibody (sheep anti mouse IgG-HRPO, 515-035-071-JIR, Stratech, UK) was added at 1/100 dilution, 50 µl per well and incubated for 1 h at 20°C. TMB substrate (50 µl per well) was added and allowed to develop for 6 min. The reaction was stopped by the addition of 100 µl 10% H_2_SO_4_ per well and plates were read at *A*_450_. A mouse monoclonal anti-Ag85b (Abcam, ab43019) used as a positive control. The mean absorbance ± SEM of each mouse group is shown. Data for each mouse, at time 0, has been normalized to the day 0 average reactivity to Ag85b in all mice screened.

## Acknowledgements

This research was funded by the Biotechnology and Biological Sciences Research Council (BBSRC Follow On Fund BB/N022165/1, awarded to JvdE and KJM). AAW was supported by a PhD scholarship granted Raoul and Catherine Hughes and the University of Bath Alumni. KJM, HD and RW were also supported by the MRC and Newcastle University’s Confidence in Concept funding. AS thanks the Royal Society URF and Alumni Fund at the University of Bath for funding. MAG was supported by the EMBL interdisciplinary Postdoc Programme under Marie Curie COFUND Actions as well as the Horizon 2020 programme of the European Union, iNEXT (H2020 Grant # 653706).

## Author Contributions

JvdE, AGW and KJM conceived the idea of the project. YY, KJM and JvdE designed the experiments. YY, CRB, AAW, RK and JP performed and analysed the experiments. KW and WK conducted and analysed the *switch* SENSE experiments. MAG and DIS conducted SAXS experiments and oversaw the structural analysis. RK, JP and KJM conducted the *in vivo* experiments. AS and AW significantly contributed to the discussions about the overall project. YY, CRB, KJM and JvdE wrote and edited the manuscript, with significant contributions from AAW, MAG and DIS.

## Data availability

All data generated or analysed during this study are included in this published article (and its supplementary information files).

